# CyTOF reveals phenotypically-distinct human blood neutrophil populations differentially correlated with melanoma stage

**DOI:** 10.1101/826644

**Authors:** Yanfang Peipei Zhu, Tobias Eggert, Daniel J. Araujo, Pandurangan Vijayanand, Christian H. Ottensmeier, Catherine C. Hedrick

**Author notes:** Corresponding Author: Catherine C. Hedrick, La Jolla Institute for Allergy and Immunology, 9420 Athena Circle, La Jolla, CA 92037.

## Abstract

Understanding neutrophil heterogeneity and its relationship to disease progression has become a recent focus of cancer research. Indeed, several studies have identified neutrophil subpopulations associated with pro- or anti-tumoral functions. However, this work has been hindered by the lack of widely-accepted markers with which to define neutrophil subpopulations. To identify markers of neutrophil heterogeneity in cancer, we utilized single-cell cytometry by time-of-flight (CyTOF) coupled with high-dimensional analysis on blood samples from treatment-naïve, melanoma patients. Our efforts allowed us to identify 7 blood-neutrophil clusters, including 2 previously identified individual populations. Interrogation of these neutrophil subpopulations revealed a positive trend between specific clusters and disease stage. Finally, we recapitulated these 7 blood-neutrophil populations via flow cytometry and found that they exhibit diverse capacities for phagocytosis and reactive oxygen species (ROS) production *in vitro*. In summary, our data provide a refined consensus on neutrophil-heterogeneity markers, enabling a prospective functional evaluation in patients with solid tumors.

**KEY POINTS:**

- CyTOF analysis reveals 7 blood neutrophil clusters correlating with melanoma stage in treatment-naïve patients.
- Neutrophil clusters by unbiased-calling are recapitulated by flow cytometry and harbor diverse phagocytic and ROS-producing capacities.

## INTRODUCTION

Neutrophils are bone marrow (BM)-derived myeloid cells that play pivotal roles in anti-cancer immunity.^1^ These cells are produced at a rate of 10^11^ per day ^2,3^ and they comprise 50 - 70% of blood leukocytes when released into circulation. Due to this rapid turn-over in the body, neutrophils have traditionally been viewed as a homogeneous population. However, recent work has shown that they exhibit a longer life cycle than previously thought ^4^; reviving interest in the possibility of distinct neutrophil populations.^3,5^

Spurred on by such findings, several groups have identified and characterized neutrophil subpopulations. For example, use of density gradient separation has uncovered distinct, low-density (LDN) and high-density neutrophil (HDN) populations, each with opposing actions in immune regulation and cancer progression.^6^ Marini and coauthors employed flow cytometry to show that CD10^+^ and CD10^-^ neutrophils represent cell populations with opposing effects on T cell proliferation.^7^ Pillay and colleagues identified 3 neutrophil subpopulations, based on their differential expression of CD16 and CD62L, which exhibit specific maturation and activation statuses.^8^ CD45RA-positive expression in certain neutrophils was also found to indicate activation statuses.^9^ Singhal and collaborators isolated a CD14^+^ neutrophil subpopulation with anti-tumor functions, including enhancement effector T cell production of IFN-g and Granzyme B.^10^ Evrard and colleagues have demonstrated a CD15^+^CD49^+^CD101^-^ neutrophil precursor (preNeu).^11^ Our group has also identified a CD117^+^CD66b^+^CD38^+^ neutrophil progenitor (NeP), which was found in the blood of tumor-bearing animals.^12^ Additional work in this area is summarized in two excellent review articles.^13,14^ Nevertheless, a lack of widely-accepted subpopulation markers has impeded our understanding of neutrophil heterogeneity. Indeed, the neutrophil subpopulations thus far described likely represent overlapping populations. Flow cytometry analysis suggests that CD10^+^ and CD10^-^ neutrophil subpopulations are fractionated into both LDN and HDN layers.^7^ Furthermore, the CD10^+^ expression demonstrated by Marini et al^7^ is shared by the CD16^bright^ subpopulation reported by Pillay et al.^8^ CD14^+^ neutrophils express CD10^-^,^10^ suggesting that this subpopulation overlaps with the CD10^-^ neutrophil population.^7^ The CD14^+^ neutrophil subpopulation also expresses CD49d^+^, indicating that it could also overlaps with the CD49d^+^ preNeu demonstrated by Evrard et al,^11^ as well as a CD49d^+^CD62L^lo^ neutrophil subpopulation (“aged neutrophil”) reported by Zhang et al.^15^ To determine the extent to which previously reported neutrophil subpopulations intersect, high-dimensional analysis of neutrophil heterogeneity on single-cell basis is imperative.

We and others have employed high-dimensional approaches such as single-cell cytometry by time of flight (CyTOF) and single-cell RNA sequencing (scRNA-seq) to address neutrophil heterogeneity. These endeavors demonstrate that neutrophil-lineage cells comprise a heterogeneous pool in mouse and human BM.^11,12^ Additionally, recent scRNA-seq analyses reveal 6 neutrophil clusters with distinct transcriptional signatures in human lung tumors but the surface markers needed to classify these populations were not identified.^16^ Interestingly, work in this field has also suggested differential involvement of neutrophil subpopulations in cancer.^17,18^ Thus, the development of neutrophil consensus markers is required for improving our understanding of neutrophil biology and its relationship to disease progression.

Here, we utilize a CyTOF panel of the most-commonly used surface markers of neutrophil maturation, activation, and function to comprehensively investigate neutrophil heterogeneity in whole blood (WB) from treatment-naïve melanoma patients. High-dimensional analysis of this dataset revealed 7 neutrophil subpopulations associated with disease stage and which are reproducible during manual gating in flow cytometry. Finally, we found that these 7 neutrophil subpopulations harbor distinctive functions, demonstrated by their differential capacities for phagocytosis and reactive oxygen species (ROS) production.

## METHODS

### Melanoma patient blood collection

Blood samples from melanoma patients who were treatment-naïve post-surgical resection were collected in EDTA-coated tubes by the Biospecimen Repository Core Facility (BRCF) at the University of Kansas Cancer Center and delivered to La Jolla Institute for Immunology (LJI) via overnight shipping. Concurrently, blood from healthy donors were collected in EDTA-coated tubes at LJI. To ensure uniform treatment between control and experimental materials, all healthy donor blood samples were stored at 4°C overnight and then processed at the next morning.

### Healthy donor blood collection

EDTA-coated blood from healthy volunteers was obtained after written informed consent under the guidelines of the Institutional Review Board of LJI and in accordance with the United States Department of Health and Human Services’ Federal Policy for the Protection of Human Research Subjects (VD-057-0217).

### Cell suspension from human whole blood

Whole blood (WB) was subject to red blood cell lysis (RBC lysis buffer, eBiosciences) twice at room temperature (RT) for 10 min. Cells were then washed with staining buffer (D-PBS + 1 % human serum + 0.1 % sodium azide + 2 mM EDTA) and filtered through a 70 μm strainer. Cell suspensions were prepared by sieving and gentle pipetting to reach final concentration of 3 x 10^6^ cells per 100 uL buffer.

### Mass cytometry antibodies

Metal-conjugated antibodies were purchased directly from Fluidigm for available targets. For all other targets, purified antibodies were purchased as described before.^19^ Antibody conjugations were prepared using the Maxpar Antibody Labeling Kit (Fluidigm) according to the manufacturer’s recommendations. Afterwards, Maxpar-conjugated antibodies were stored in PBS-based antibody stabilization solution (Candor Biosciences) supplemented with 0.05% sodium azide at 4°C. All antibodies were titrated before use.

### Mass cytometry (CyTOF)

CyTOF was performed following previously described protocols.^20^ For viability staining, cells were washed in PBS and stained with Cisplatin (Fluidigm) at a final concentration of 5 μM. Prior to surface staining, RBC-lysed WB cells were resuspended in staining buffer for 15 minutes at RT to block Fc receptors. The surface antibody cocktail listed in Table 2 was added into cell suspensions for 1 hour at 4°C. The cells were then washed with staining buffer and fixed with 1.6 % paraformaldehyde (Thermo Fisher) for 15 minutes at RT. Afterwards, 1 mL of intercalation solution for each sample was prepared by adding Cell-ID Intercalator-Ir (Fluidigm) into Maxpar Fix and Perm Buffer (Fluidigm) to a final concentration of 125 nM (a 1,000X dilution of the 125 μM stock solution) and vortex to mix. After fixation, the cells were resuspended with the intercalation solution and incubated overnight at 4°C. Cells were then washed in staining buffer, and then with subsequent washes in Cell Acquisition Solution (CAS) (Fluidigm), to remove buffer salts. Next, the cells were resuspended in CAS with a 1:10 dilution of EQ Four Element Calibration beads (Fluidigm) and filtered through a 35 μm nylon mesh filter cap (Corning, Falcon). Samples were analyzed on a Helios 2 CyTOF Mass Cytometer (Fluidigm) equipped with a Super Sampler (Victorian Airship & Scientific Apparatus) at an event rate ≤ 500 events/second. Mass cytometry data files were normalized using the bead-based Normalizer ^21^ and analyzed using Cytobank analysis software (https://www.cytobank.org/). For FLOWSOM analysis in Cytobank, hierarchical clustering was used to determine seven meta-clusters based on median markers expression (after arcsinh transformation with cofactor equal to 5) from the viSNE results.

### Flow cytometry and cell sorting

All FACS staining was performed in staining buffer at 4°C. Cells were filtered through sterile, 70 μm cell strainers to obtain single-cell suspensions (30,000 cells per μL for flow cytometry analysis, 0.5 - 2 x 10^7^ cells per mL for sorting). Prior to surface staining, RBC-lysed WB cells were resuspended in staining buffer to block Fc receptors for 15 minutes at RT. Surface staining was performed for 30 minutes in a final volume of 500 uL for FACS sorts and 100 uL for standard flow cytometry. Cells were washed twice in at least 200 uL FACS buffer before acquisition. FACS was performed via an Aria II and Aria-Fusion (BD biosciences) and conventional flow cytometry via an LSRII and LSR Fortessa (BD Biosciences). All flow cytometry was performed on live cells. Percentages of CD45^+^ immune cells were calculated by forward and side scatter and viability analyses of live cells. All analyses and sorts were repeated at least 3 times, and the purity of each sorted fraction was determined visually and by FACS reanalysis of surface markers. Data were analyzed using FlowJo software (version 10.5).

### Phagocytosis

Phagocytic capacities of neutrophils were assessed with a Phagocytosis Assay Kit (Red Zymosan) (Abcam) following the manufacturer’s protocol. RBC-lysed WB cells were diluted to a concentration of 3 x 10^6^ cells in 1 mL buffer and incubated with 5 μL of Zymosan slurry per sample at 37°C and 5 % CO_2_ for 2.5 hours. Afterwards, the samples were centrifuged for 5 minutes at 400 x g and then stained with 100uL antibody cocktail for flow cytometry-based detection of phagocytosis of Red Zymosan particles by neutrophils.

### Cellular ROS detection

ROS production by neutrophils was detected with a Cellular ROS Detection Assay Kit (Abcam) following manufacturer’s manual. The RBC-lysed WB cells were diluted to the concentration of 3 x 10^5^ cells in 100 mL buffer. Pre-treatment with ROS inhibitor (N-acetyl-L-cysteine) was carried out for the negative control group at 37°C, 5 % CO_2_ for 30 minutes. ROS detection antibody then added into the antibody cocktail for flow cytometry-based detection of neutrophil ROS production. ROS inducer (pyocyanin) was added to all groups and incubated for 30 minutes prior to acquisition on the cytometer.

### Quantification and statistical analyses

Data for all experiments were analyzed with Prism software (GraphPad). For Figure 2, linear regression and Pearson correlation coefficients were used to determine the correlation between neutrophil cluster frequencies with melanoma stages. P values were calculated based on twotailed comparisons with 95 % confidence intervals and shown in APA style. For all bar graphs, error bars indicate mean ± SD.

## RESULTS

### CyTOF reveals 7 neutrophil clusters in blood from melanoma patients

In order to identify novel blood neutrophil populations, we utilized cytometry by time of flight (CyTOF) to analyze red blood cell (RBC)-lysed fresh blood samples from a cohort of 21 melanoma patients (Table 1). At the time of sample collection, these patients were recently diagnosed, had not received any treatment for their condition post-surgery (treatment-naïve), and ranged from 24 to 82 years of age, with a median age of 69 years (Table 1). After processing, samples were subjected to a CyTOF antibody panel that simultaneously measured the expression of 40 neutrophil surface markers (Table 2). Leukocyte lineage markers (Table 2) were used to perform visualization of t-distributed stochastic neighbor embedding (viSNE)-automated analysis to study live CD45^+^ single cells in blood (Supplemental Figure 1A). Neutrophils (CD66b^+^ cell-enriched cluster) were distinguishable from all other leukocytes using this strategy (Supplemental Figure 1B).

**Table 1.** Patient demographics and tumor characteristics.

Characteristics
|  |  |  |
| --- | --- | --- |
| Age, y | Median (range) | 69 (24-82) |
| Sex, No. (%) |  |  |
|  | Male | 12 (57.1) |
|  | Female | 9 (42.9) |
| Primary tumor thickness, mm |  |  |
|  | No. (range) | 13 (0.4 - 16) |
|  | 0–1.00, No. (%) | 4 (19) |
|  | 1.00–4.00, No. (%) | 6 (28.6) |
|  | >4.00, No. (%) | 3 (14.3) |
|  | Missing, No. (%) | 8 (38.1) |
| Primary tumor ulceration status, No. (%) |  |  |
|  | Absent | 9 (42.9) |
|  | Present | 4 (19) |
|  | Missing | 8 (38.1) |
| Primary tumor anatomic site, No. (%) |  |  |
|  | arms/legs | 5 (23.8) |
|  | torso | 5 (23.8) |
|  | head/neck | 7 (33.3) |
|  | Missing | 4 (19) |
| Primary tumor mitosis, No. (%) |  |  |
|  | Absent | 5 (23.8) |
|  | Present | 7 (33.3) |
|  | Missing | 9 (42.9) |
| AJCC* stage at pathological diagnosis, No. (%) |  |  |
|  | Stage I | 8 (38.1) |
|  | Stage II | 4 (19) |
|  | Stage III/IV | 5 (23.8) |
|  | Missing | 4 (19) |

**Table 2.** CyTOF antibody panel.

| Leukocyte Lineage |  |  | Maturation |  |  | Heterogeneity (function) |  |  |
| --- | --- | --- | --- | --- | --- | --- | --- | --- |
| Isotope | Metal | Specificity | Isotope | Metal | Specificity | Isotope | Metal | Specificity |
| 89 | Y | CD45 | 146 | Nd | CD64 | 153 | Eu | CD14 |
| 113 | In | CD3 | 151 | Eu | CD49d | 161 | Dy | Arg1 |
| 113 | In | CD127 | 154 | Sm | CD117 | 168 | Er | CD304 (Nrp1) |
| 115 | In | CD41 | 156 | Gd | CD10 | 171 | Yb | HLA-A/B/C |
| 115 | In | CD235a | 158 | Gd | CD101 | 174 | Yb | HLA-DR |
| 141 | Pr | CD11c | 165 | Ho | CD16 | Heterogeneity (migration) |  |  |
| 143 | Nd | CD123 | 166 | Er | CD34 | Isotope | Metal | Specificity |
| 143 | Nd | CD203c | 167 | Er | CD38 | 147 | Sm | CD182 (CXCR2) |
| 144 | Nd | CD19 | 172 | Yb | CD15 | 159 | Tb | CD197 (CCR7) |
| 148 | Nd | CD11b | Heterogeneity (adhesion/ activation) |  |  | 175 | Lu | CD184 (CXCR4) |
| 152 | Sm | CD66b | Isotope | Metal | Specificity | Proliferation |  |  |
| 163 | Dy | CD86 | 142 | Nd | CD11a (LFA-1) | Isotope | Metal | Specificity |
| 164 | Dy | Siglec 8 | 145 | Nd | CD62L | 127 | I | IdU |
| 169 | Tm | CD33 | 149 | Sm | CD48 | 162 | Dy | CD71 |
| 176 | Yb | CD56 | 155 | Gd | CD45RA |  |  |  |
| Heterogeneity (other) |  |  | 170 | Er | CD35 |  |  |  |
| Isotope | Metal | Specificity |  |  |  |  |  |  |
| 160 | Gd | CD79b |  |  |  |  |  |  |

We focused on the CD66b^+^ cell-enriched cluster to analyze the heterogeneity of neutrophils across all patient samples. Our results demonstrate that these samples contained a CD117^+^CD66b^+^, hNeP population (Figure 1A), which we formerly identified.^12^ In addition, FlowSOM analyses^22^ revealed another 6 neutrophil clusters (termed N_1_ through N_6_, Figure 1B), each with distinct surface marker profiles (Figure 1C). As maturity markers are commonly used to distinguish circulating neutrophil subpopulations in cancer^11,23^, we next quantified expression levels of these markers in the N_1_-N_6_ clusters. We found that cluster N_2_ expresses high levels of CD101, CD10, and CD16 compared with the other clusters, indicating that this cluster represents terminally differentiated, mature neutrophils (Figure 1C and D).^7^ Furthermore, CD10^lo^ clusters (N_1_, N_3_-N_6_) express reduced levels of CD101, highlighting their immaturity when compared to the N_2_ cluster (Figure 1D). The N_1_, N_3_, and N_6_ clusters express progenitor markers (CD34 and CD117) at a lower level than hNeP, suggesting a precursor phenotype for these 3 clusters (Figure 1D; supplemental Figure 2A). In the N_5_ cluster, expression of CD34 and CD117 was diminished, compared to the N_1_, N_3_, and N_6_ clusters. In contrast, CD10 and CD16 levels were higher in the N_5_ cluster, compared to N_3_ and N_6_, intimating that this cluster denotes nonproliferative immature neutrophils (Figure 1D). Moreover, the N_3_ and N_6_ neutrophil clusters both express CD101^-^CD49d^+^, which suggests these clusters belong to the neutrophil precursor (preNeu) population.^11^

**Figure 1.**
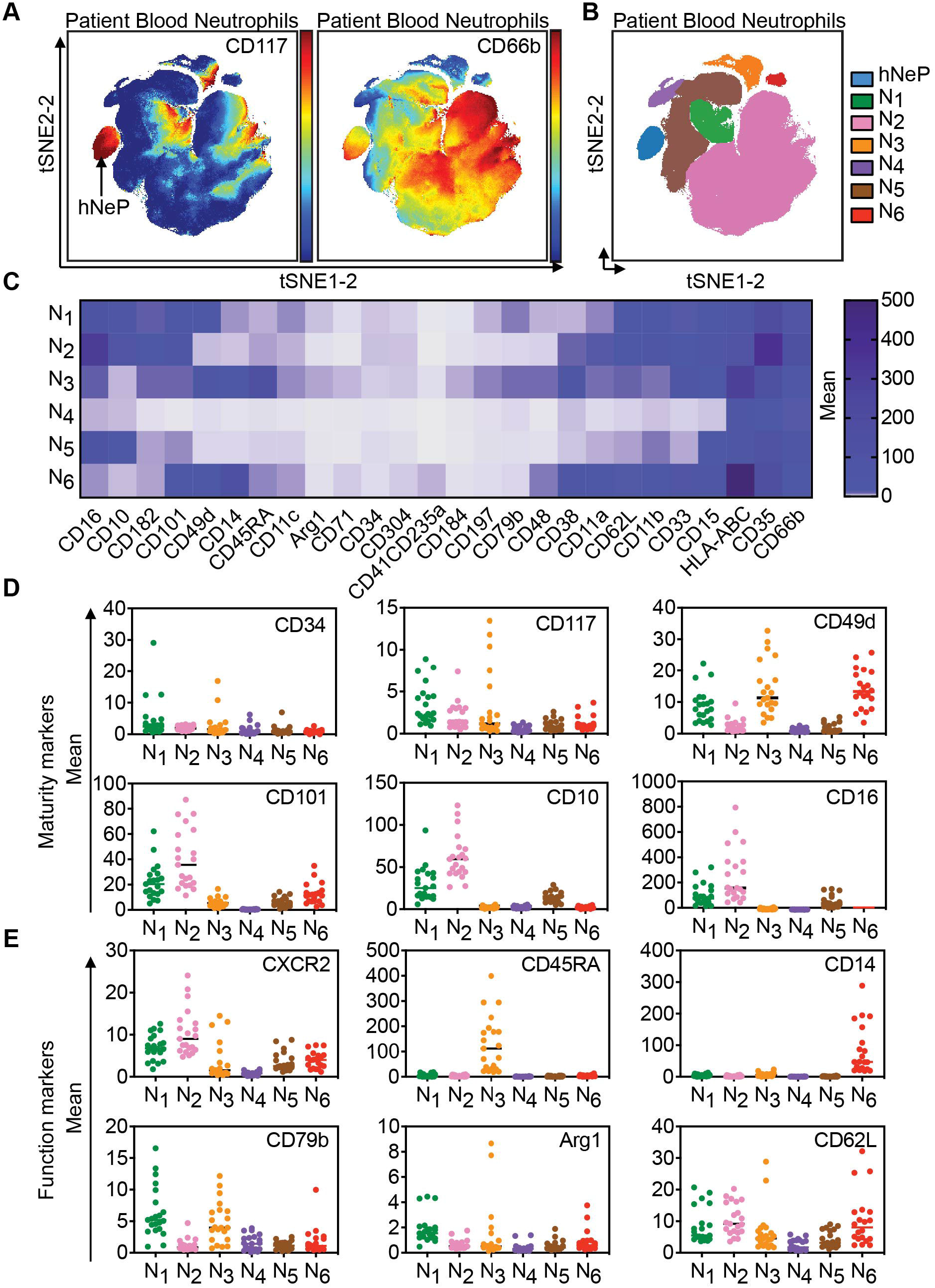
CyTOF-based analysis of blood from melanoma patients reveals 7 automated neutrophil clusters. The treatment-naïve melanoma patients’ blood neutrophils (the CD66b^+^ automated cluster from supplemental Figure 1B) were subjected to automated analysis. (A) Mean intensities of CD117 and CD66b expression are shown on viSNE map as spectrum colored dots (low in blue, high in red). hNeP was identified on the viSNE map based on the expression of CD117^+^CD66b^+^. (B) FLOWSOM analysis of the viSNE results revealed 7 automated clusters. (C) Heatmap shows the mean intensity of each marker in the 6 unidentified automated clusters on a global scale. (D) Dot plot shows the mean intensity of each maturity marker in the 6 unidentified automated clusters. Each dot represents result of one patient. (E) Dot plot shows the mean intensity of each functional marker in the 6 unidentified automated clusters. Each dot represents result of one patient.

We further characterized these neutrophil clusters by comparing functional markers previously used to evaluate neutrophil populations (Figure 1E). In agreement with initial findings (Figure 1C), N_2_ showed the highest CXCR2 levels, confirming its maturation status. Interestingly, N_3_ and N_6_, which are both from the CD101^-^CD49d^+^ preNeu pool, expressed different levels of CD45RA and CD14. CD45RA was expressed only by N_3_ whereas CD14 was exclusively expressed by N_6_. Additionally, N_3_ exhibited higher CXCR4 levels, compared with cluster N_6_, which categorizes this group as likely the previously reported, “aged” or senescent (CXCR4^+^CD49d^+^CD62L^lo^) neutrophil population (supplemental Figure 2B).^15^ Finally, N_1_ and N_6_ expressed low amounts of CD16, compared to N_2_ (Figure 1D), and stained positive for CD62L (Figure 1E), suggesting these cells are the CD16^dim^CD62L^bright^ band cells that have been described prior.^8^ Overall, our approach is able to call both novel and established neutrophil populations from melanoma patients.

### Neutrophil heterogeneity correlates with melanoma stage

Based on the aforementioned results, we sought to determine the frequency of each neutrophil cluster in our patient samples. We did not observe significant correlations between specific clusters or patient demographics such as age and sex, nor tumor characteristics such as anatomice site or ulceration status (Table 1). However, the overall frequency of precursors/immature neutrophils (hNeP, N_1_, N_3-5_) increased up to 4-fold with disease prognosis (Figure 2A, supplemental Figure 3A). Regression analysis of each cluster’s (hNeP and N1-N6) percentage within total neutrophils determined that decreased N_2_ frequencies correlated with melanoma stage (Pearson r = 0.5473, p = 0.023) (Figure 2B). Thus, we hypothesized that different patterns of neutrophil heterogeneity demonstrated by cluster frequencies are predictive of disease stage. To test this hypothesis, we categorized patients into 4 different groups (A-D) based on these patterns, as analyzed by viSNE (Figure 2C, supplemental Figures 3B and C). From this analysis, we determined that patients in Group C have the highest hNeP, N_1_ and N_3_ frequencies. Patients included in Group D have the highest N_4_ and N_5_ frequencies (Figure 2D). The percentage of the mature, N_2_ cluster progressively decreases from patient Group A to D. We next investigated the melanoma stage of the patients in these 4 groups (Figure 2E). Interestingly, patients in Group A were all diagnosed with early stage cancer whereas groups C and D contained the highest percentages of stage III and stage IV patients. These results suggest that specific patterns of neutrophil heterogeneity are associated with melanoma progression and may assist in patient grouping and diagnosis.

**Figure 2.**
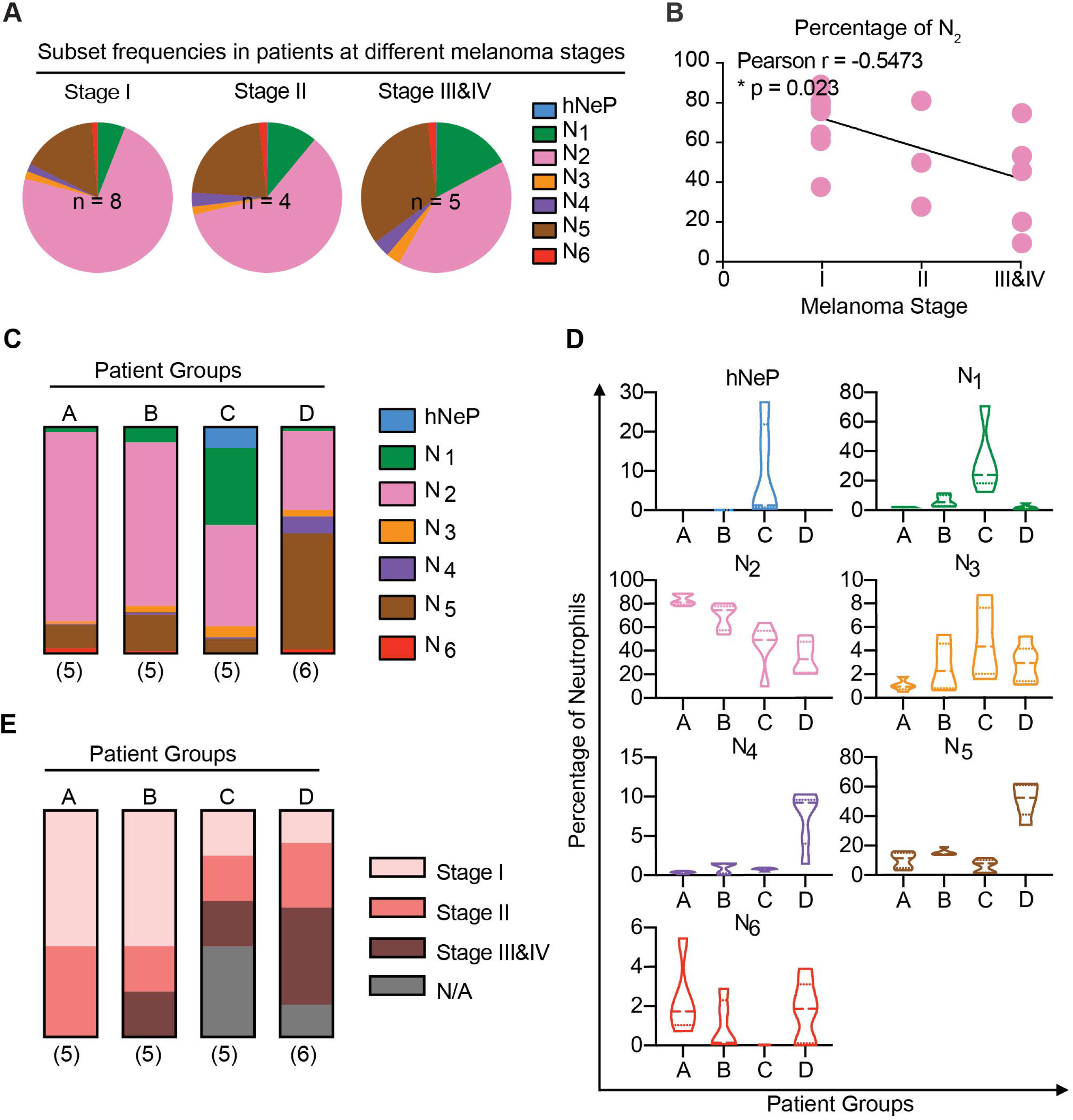
Neutrophil heterogeneity in melanoma patients correlates with disease stage. (A) Line regression analysis shown in dot plots depicting correlations between neutrophil cluster frequencies and melanoma stage. Each dot represents one patient. Pearson analysis results are shown for each cluster. P values were calculated based on two-tailed comparisons with 95 % confidence intervals and shown in APA style. (B) Pie charts show mean percentages for each FLOWSOM cluster (hNeP and N_1_-N_6_) in total blood neutrophils of patients grouped by melanoma stage. Only patients with a melanoma stage diagnosis shown in Table 1 were used for this analysis. The numbers of subjects in each melanoma stage are indicated on the graph. (C) Bar graph shows the mean percentage of each cluster in patient Groups A-D. All patients were used for this analysis, regardless of whether or not they received a melanoma stage diagnosis (Table 1). The numbers of subjects in each group are indicated below each column. (D) Violin plots shows the neutrophil cluster frequency in patient pools A-D. Quartiles and median value of each patient pool are indicated as dotted lines. All patients were used for this analysis, regardless of whether or not they received a melanoma stage diagnosis (Table 1). (E) Bar graph shows the percentage of patients at different melanoma stage (Table 1) in patient Groups A-D. N/A indicates missing diagnosis information of patients. All the patients were used for this analysis, regardless of whether or not they received a melanoma stage diagnosis (Table 1). The numbers of patients in each pool are indicated below each column.

### Flow cytometry recaptures the 7 neutrophil subpopulations in melanoma patients’ blood

We then asked if the markers highlighted in our *in silico* analysis could be used to devise a manual gating strategy for replicating the 7 neutrophil clusters identified by CyTOF (Figure 3A; supplemental Figure 4). Thus, we first used manual gating to isolate total neutrophils from the CyTOF data. These cells overlapped with 90% of our automated CD66b^+^ cell-enriched cluster by backgating (Supplemental Figure 4). By first applying a manual gate for CD66^+^ neutrophils, we were able to identify the 7 subpopulations by their expression patterns of CD117, CD79b, CD45RA, CD16, CD49d, CD101, and CD10. To verify whether manually gated populations mirror automated clusters, we validated this gating strategy by: 1) back-gating the manually-gated subpopulations to the automated viSNE map and 2) back-gating automated clusters to the manual gates (supplemental Figure 5A, B). Both methods confirmed that our manual gating strategy is able to successfully isolate the CyTOF-identified hNeP and N_1_-N_6_ clusters.

**Figure 3.**
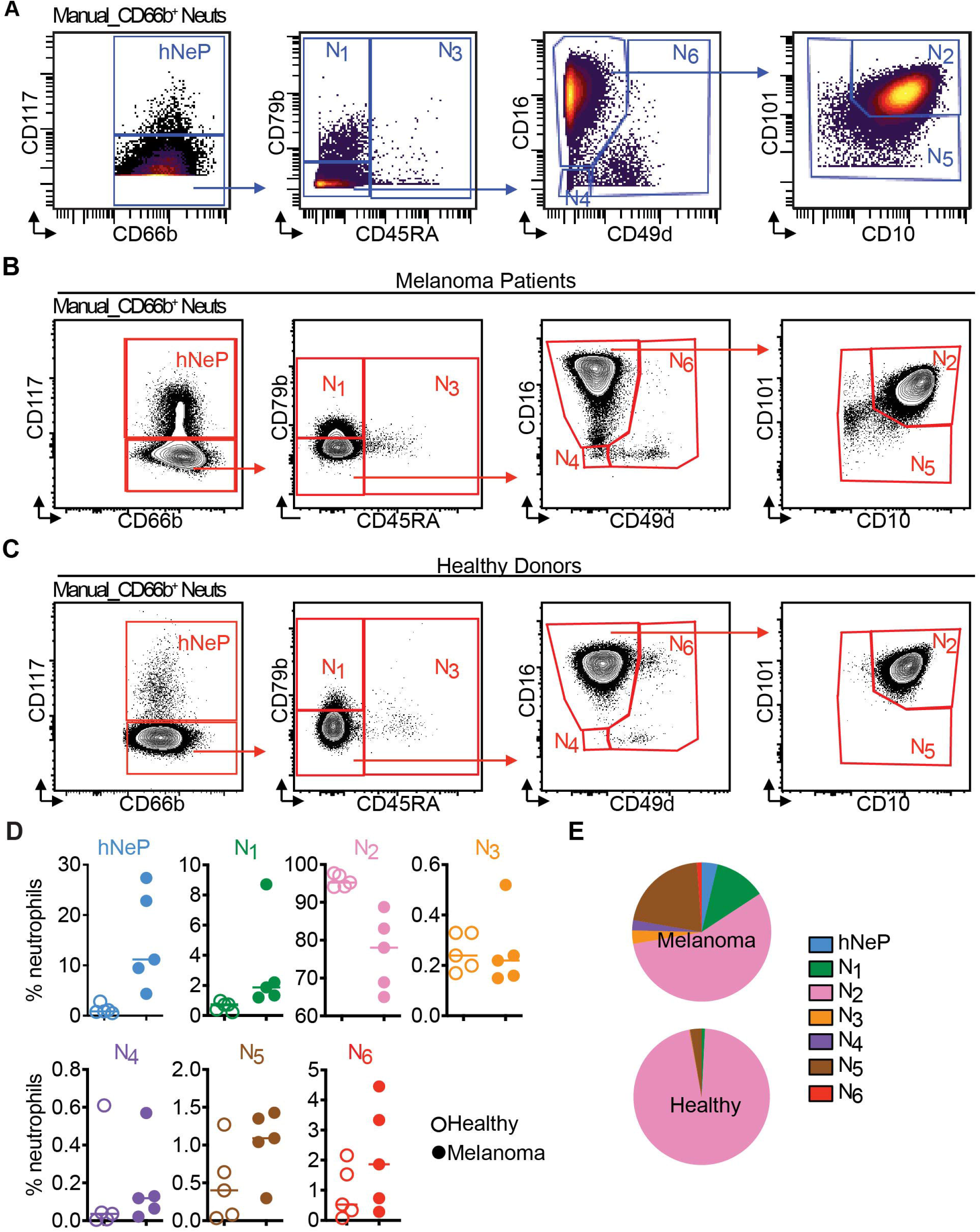
Flow cytometry replicates the 7 neutrophil subpopulations. (A) CD66b^+^ blood neutrophils were manually selected and subjected to sequential gating to identify the neutrophil subpopulations with CyTOF. Scales are shown in arcsinh transformation with cofactor equal to 5. (B) The gating strategy from (A) was validated by flow cytometry in treatment-naïve melanoma patients. Scales are shown in biexponential scale. (C) Flow cytometry analysis of healthy donor’s blood neutrophils with the gating strategy from (A). Scales are shown in biexponential scale. (D) Flow cytometry analysis of the frequency of each manually gated neutrophil subpopulation in total blood neutrophils. 5 healthy donors (age 23-46, 2 female, 3 male) and 5 treatment-naïve melanoma patients (aged 59-79 years, 2 female and 3 male) were analyzed. Each dot represents result of one patient. Scales are shown in biexponential scale. (E) Pie charts show the mean percentage of each cluster based on the viSNE analysis of healthy donor’s blood neutrophils compared to treatment-naïve melanoma patients from Figure 1. 2 healthy donors were analyzed.

Next, we further validated this manual gating strategy (Supplemental Figure 4; Figure 3A) via flow cytometry (Supplemental Figure 5C; Figure 3B, C). To validate the utility of these findings as they relate to cancer, we investigated neutrophil heterogeneity in 5 healthy donors (age 23-46, 2 female, 3 male) and compared them to 5 additional treatment-naïve melanoma patients (aged 59-79 years, 2 female and 3 male). We observed that the N_2_ population comprises >95% of total neutrophils in healthy donors, whereas the other populations (hNeP, N_1_, N_3_, N_4_, N_5_, N_6_) were rarely detected (Figure 3C, D). Moreover, compared to healthy donors, these subpopulations (hNeP, N_1_, N_3_, N_4_, N_5_, N_6_) appeared more frequently in treatment-naïve melanoma patients, which occupied >10% total neutrophils, as determined by flow cytometry. The frequencies of N_2_, however, were reduced to <90% of total neutrophils in melanoma-patient blood. Finally, we randomly selected blood from two of these additional healthy controls and performed CyTOF with the same high-dimensional analysis settings that we used to generate the results in Figure 1 (Supplemental Figure 6). Confirming our results above, neutrophils from healthy donors contained over 95% of the mature, N_2_ subpopulation (Figure 3E). This result is consistent with prior findings showing that both immature neutrophils and neutrophil precursors are mobilized to the circulation in cancer. ^24,25^

### The 7 neutrophil subpopulations perform diverse immune functions

Afterwards, we sought to determine whether the 7 neutrophil subpopulations we identified express different immunological phenotypes. Phagocytosis and generation of reactive oxygen species (ROS) are two major functions of neutrophils in the immune system.^26^ Therefore, we evaluated the 7 neutrophil subpopulations’ phagocytic and ROS-producing capacities.

The 7 neutrophil subpopulations from 3 randomly selected melanoma patients (Age 59, 77, 79, 1 female and 2 males) were analyzed for their phagocytosis of pre-labeled Zymosan particles. Uptake of Zymosan particles by neutrophil subpopulations were then determined by gating with flow cytometry (red in the +Zymosan group, Figure 4A top panels), and compared with the control-Zymosan group (Figure 4A bottom panels). To determine each subpopulation’s phagocytic ability, we compared the Zymosan positive portion (Zym+ Neuts) in each neutrophil subpopulation from the +Zymosan group. Strikingly, each neutrophil subpopulation performed phagocytosis at different levels of efficiency. Our results show that the hNeP and N_1_ populations display the highest phagocytic capacities (Figure 4B, C). Indeed, the phagocytic abilities of hNeP and N_1_ were comparable to, or higher than, that of monocytes collected from the same donors (supplemental Figure 7A, B). Moreover, the phagocytic capabilities of the other subpopulations were reduced by 3- to >20-fold (Figure 4B, C). Despite the fact that N_2_ and N_5_ harbor lower phagocytic capacity compared to hNeP and N_1_, they are the most prevalent neutrophil subpopulations in melanoma patients’ blood (Figure 1 and 2). Therefore, the absolute numbers of Zym+ N_2_ and N_5_ cells still comprise the largest portion (about 50%) of all phagocytic (Zym+) neutrophils and about 25% of total phagocytic CD45^+^ leukocytes (supplemental Figure 7C, D).

**Figure 4.**
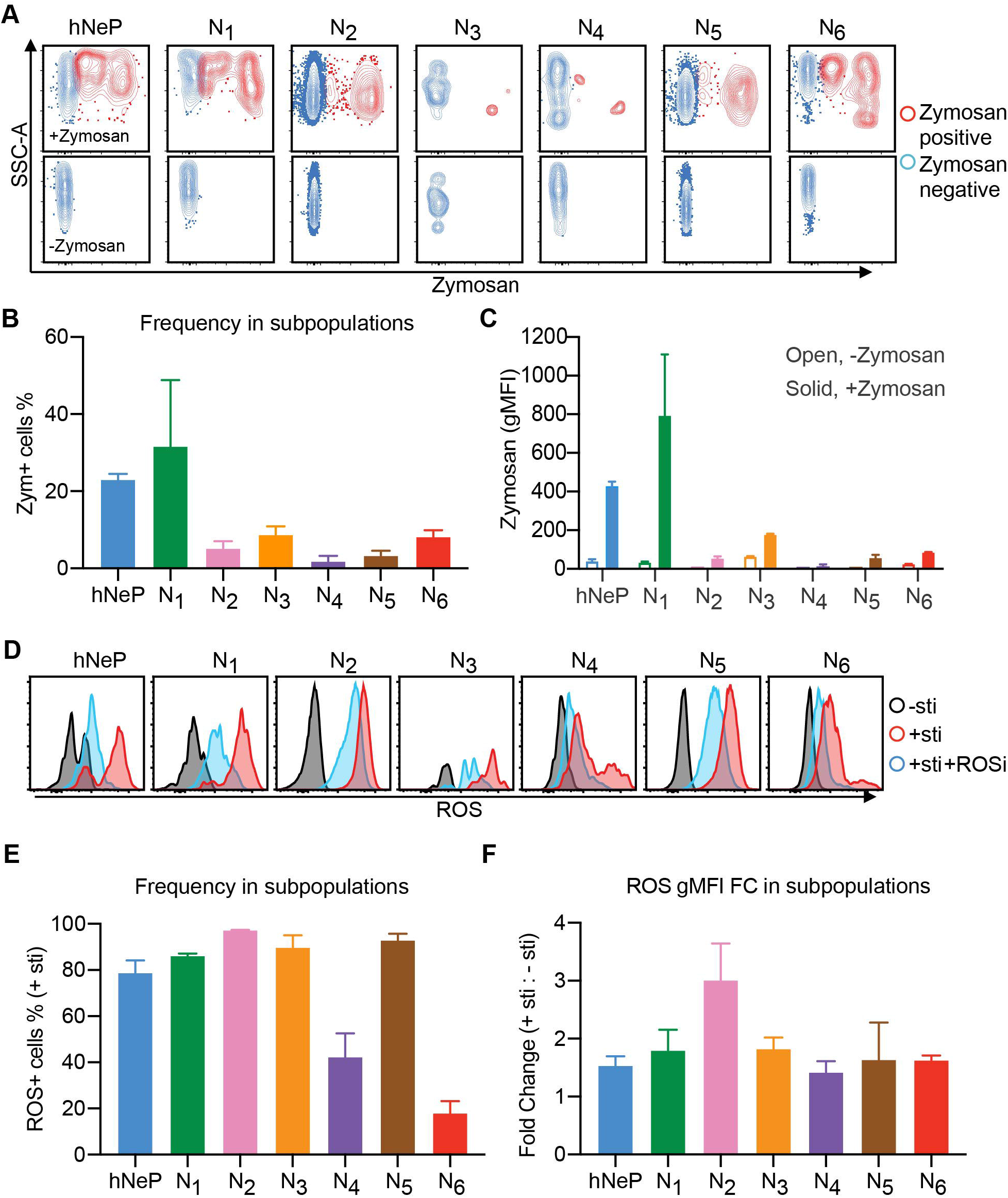
The 7 neutrophil subpopulations harbor diverse phagocytic and ROS producing capacities. 3 randomly selected melanoma-naïve patients (Age 59, 77, 79, 1 female and 2 males) were analyzed with flow cytometry. (A-C) RBC-lysed blood samples were incubated with pre-labeled Zymosan particles for 2.5 hours. Afterwards, the cells were harvested and stained with the flow cytometry panels described in Figure 3 and supplemental Figure 5C. Each neutrophil subpopulation was gated to evaluate its uptake of Zymosan particles. (A) The Zymosan positive cells are shown in red, the Zymosan negative cells are shown in blue. The no zymosan group (-Zymosan) is shown in bottom panels as the control group. (B) The percentage of the Zym+ cells (red dots in (A)) in each gated neutrophil subpopulation. Error bars indicate mean with SD. (C) The geometric MFI of Zymosan in each gated neutrophil subpopulation. Error bars indicate mean with SD. (D) The 7 neutrophil subpopulations’ ability to produce ROS is determined by flow cytometry. RBC-lysed blood samples were split into 3 groups and incubated with: no stimulation (- sti), ROS inducer (Pyocyanin) alone (+ sti), or ROS inducer plus ROS inhibitor (+ sti + ROSi). Afterwards, each neutrophil subpopulation was gated for evaluation of ROS+ cells. Histogram plots show the gated neutrophil subpopulations in each group: - sti is shown in black, + sti is shown in red, + sti + ROSi is shown in blue. (E) The percentage of the ROS+ cells in each gated neutrophil subpopulation from the + sti group. Error bars indicate mean with SD. (F) The ROS geometric MFI fold change (FC) of each gated neutrophil subpopulation from the + sti group to the - sti group. Error bars indicate mean with SD.

Finally, we sought to determine the ability of neutrophil subpopulations to produce ROS. The 7 neutrophil subpopulations from 3 randomly selected melanoma-naïve patients (Age 59, 77, 79, 1 female and 2 males) were analyzed by flow cytometry. The ability to produce ROS was evaluated in 3 experimental groups: no stimulation (- sti), ROS inducer (Pyocyanin) alone (+ sti), or ROS inducer plus ROS inhibitor (+ sti + ROSi). The production of ROS was determined in gated neutrophil subpopulations. We ascertained that all 7 neutrophil subpopulations produced ROS upon stimulation with inducer and responded to inhibitor (Figure 4D). However, these cells significantly differ from one another in terms of their ability to produce ROS. For instance, the N_2_ and N_5_ displayed the highest capacity to produce ROS as determined by the percentage of ROS+ cells in the inducer alone (+ sti) group (Figure 4E). Meanwhile, populations N_4_ and N_6_ showed the lowest ROS-producing capacity of all the subpopulations, with levels comparable to those produced by monocytes (Figure 4E; supplemental Figure 7E). To eliminate confounding by non-inducer-specific ROS production by each population during incubation, we compared the fold-change (FC) of ROS gMFI in the inducer alone (+ sti) group to the no stimulation (- sti) group. The FC result agreed with our observation that the N_2_ has the highest ROS-producing capacity (Figure 4F). Additionally, we found that all these subpopulations respond to the ROSi at different levels. In brief, pre-treatment with ROSi blocked ROS production in N_2_ by 85% whereas the blocking efficacy reached only around 25% in N_4_ (Supplemental Figure 7F). Together, our data indicate that the 7 neutrophil subpopulations harbor different immune capabilities.

## DISCUSSION

Neutrophil heterogeneity has become an active research area, particularly with regards to cancer progression.^17^ Comprehensive examination by flow cytometry has revealed that neutrophils express a complex surface antigen profile.^27^ However, due to the limitations imposed by flow cytometry, researchers have had to arbitrarily select only a few surface markers to examine at a particular time. For example, studies in lung cancer research use one set of markers to identify their neutrophils of interest whereas studies in infectious diseases use another set of markers; this is also the case for work on the role of neutrophils in hematopoiesis and angiogenesis.^14,18^ Consequently, neutrophil subpopulations reported by different groups mostly share individual features such as CD66b and/or CD15 but vary in regards to the assessement of other surface antigens. Thus, a lack of consensus markers for neutrophils and their functional subpopulations remains a challenge in the identification of particular disease-relevant subpopulations.

High-dimensional analysis of neutrophils at the single-cell level is needed to resolve issues associated with identifying neutrophil consensus markers. To meet this demand, we and others have utilized CyTOF to investigate novel neutrophil progenitors/precursors in BM.^11,12^ Zilionis and colleagues have reported a comprehensive analysis of myeloid cell heterogeneity via single-cell transcriptomics in non-small-cell lung cancer (NSCLC).^16^ This study found 6 blood neutrophil populations with unique gene signatures in these patients, a myeloid precursor-like population was also found but remained uncharacterized. However, the genes that encode surface markers were not discussed in this study, making it a challenge to compare these neutrophil and myeloid precursors to established neutrophil subpopulations.

Here, we have designed a CyTOF panel that reflects our integration of markers reported on in the published data and which evaluates the most commonly used surface antigens. Our aim was to establish a paradigm for characterizing neutrophil heterogeneity, focusing on patients with melanoma. We have identified 7 neutrophil clusters in treatment-naïve melanoma blood by CyTOF. Each patient exhibited a distinct neutrophil heterogeneity pattern, based on each cluster’s frequency as a proportion of total neutrophils. We have identified groups of patients based on similarities between these neutrophil heterogeneity patterns. The patient groups displayed different melanoma stage categories, suggesting a link of neutrophil heterogeneity to stage of the disease, yet prospective validation is needed. We have also shown that these automated neutrophil clusters can be reproduced by manual gating in conventional flow cytometry. Furthermore, we demonstrate that these neutrophil subpopulations exhibit significantly different capacities for phagocytosis and ROS production. Thus, we anticipate that these subpopulations may play different roles in cancer initiation and/or progression.

However, it must be emphasized that the CyTOF panel used in this study is selective for human blood neutrophil- and cancer-markers and therefore likely will not capture neutrophil heterogeneity across species, organs, and/or different diseases. We also cannot rule out higher dimensions of heterogeneity within our reported subpopulations and such investigations should be the focus of future work. Furthermore, we are unable to rule out the possibility of phenotype switching between these 7 subpopulations and thus consider this notion an interesting direction for future investigation.^6,28^ A final caveat to our gating strategy is that some rare neutrophil subpopulations, such as those expressing TCR-like receptor,^29^ could be lost.

The results presented here suggest that some previously identified neutrophil subpopulations overlap with one another ^6,11,15^ and/or represent a mixed pool.^8,30^ This study has provided a flow cytometry gating strategy based on CyTOF analysis to isolate 7 distinctive neutrophil subpopulations. Our results show that these subpopulations display differing capacities to phagocytize debris and produce ROS. Interestingly, a decrease in the ability of T cells to bind pMHC and to respond to specific peptides is induced by myeloid cell-based production of ROS.^31^ Conversely, myeloid production of ROS has been shown to be indispensable for antigen-specific responsiveness in both CD4+ and CD8+ T cells.^32^ This controversial role of ROS in regulating T cell responses could be partially explained by the dual role of ROS; at low to moderate concentrations ROS is beneficial for cell survival, whereas high levels of ROS can induce cell death.^33^ Therefore, the 7 differing abilities of these neutrophil subpopulations to produce ROS suggests that they carry out different roles in regulating T cell responses in cancer.

## ACKNOWLEDGEMENTS

We would like to thank L. Padgett for helpful discussions about this manuscript, R. Wu for technical assistance, and the LJI Flow Cytometry Core for assistance with FACS sorting. The mass cytometry core was supported by NIH grant 1S10OD018499-01 (to C.K. and S.C.). The melanoma patient study was supported by the National Cancer Institute Cancer Center Support Grant P30 CA168524 and used the Biospecimen Shared Resource. This work was supported by NIH grants R01HL134236, P01HL136275, and R01CA202987 (all to C.C.H) and ADA7-12-MN-31 (04) (to C.C.H. and Y.P.Z).

## AUTHORSHIP

C.C.H. and Y.P.Z. conceptualized this project. C.C.H., Y.P.Z., and D.J.A. wrote the manuscript. Y.P.Z. designed and performed the experiments. Y.P.Z. and D.J.A. designed the figures. Y.P.Z., H.Q.D., T.E., and M.M. analyzed the data. C.O. and P.V. categorized the melanoma patient groups and contributed to Table 1 and Figure 2. All authors reviewed and edited this manuscript.

## SUPPLEMENTAL FIGURES

**Supplemental Figure 1. Gating strategies to identify the CD66b^+^ blood neutrophil cluster in melanoma patients with CyTOF.** This supplemental figure is associated with Figure 1 and Figure 4. RBC-lysed WB of the treatment-naïve melanoma patients in Table 1 were stained with the CyTOF panel in Table 2. (A) Live CD45^+^ cells were gated to determine the singlets for subsequent analysis. (B) Singlets were subjected to automated analysis. Left, mean intensities of each indicated marker were shown on viSNE map as spectrum colored dots (low in blue, high in red). Right, the blood cell types of each automated cluster were identified based on the hallmark marker expression shown at left panel and was displayed on viSNE map as dot overlays. Neutrophils (Neuts) were identified as the CD66b^+^ cluster (blue cluster).

**Supplemental Figure 2. Expression levels of markers in 7 automated blood neutrophil clusters.** This supplemental figure is associated with Figure 1. (A) Dot plot shows the mean intensity of CD117 and CD34 in the 6 unidentified automated clusters compared to hNeP. Each dot represents result of one patient. (B) Dot plot shows the mean intensity of CXCR4 in N_3_ and N_6_. Each dot represents result of one patient.

**Supplemental Figure 3. FLOWSOM clusters (hNeP and N_1_-N_6_) shown on viSNE maps categorize melanoma patients into different groups.** This supplemental figure is associated with Figure 2. (A) This figure is associated with Figure 2A and B. Pie charts show fold changes (FC) for each cluster’s mean frequency in patients grouped by melanoma stage. Only patients with a melanoma stage diagnosis (Table 1) were used for this analysis. (B) This figure is associated with Figure 2C-E. ViSNE maps show the FLOWSOM clusters (hNeP and N_1_-N_6_) in the total blood neutrophils of each patient. The patients were distributed into 4 groups (Group AD) based on similarities of the viSNE maps. (C) Pie charts show the percentage of each FLOWSOM cluster (hNeP and N_1_-N_6_) in the total blood neutrophils of patient groups A-D.

**Supplemental Figure 4. Manual gating strategy to select CD66b^+^ blood neutrophils with CyTOF.** This supplemental figure is associated with Figure 3A and supplemental Figure 1A. Singlets gated in supplemental Figure 1A were subjected to manual gating strategy. CD66b^+^ blood neutrophils were manually selected by sequential gating, backgated to the viSNE map (green), and then overlapped with automated CD66b^+^ cluster (blue). Scales are shown in arcsinh transformation with cofactor equal to 5.

**Supplemental Figure 5. Manual gating strategy recapitulates automated neutrophil clusters.** This supplemental figure is associated with Figure 1B and Figure 3A. (A) Automated neutrophil clusters shown in Figure 1B (names of clusters are indicated on top of each plot) were back gated to the manual gates shown in Figure 3A. The corresponding manual gates of in each plot are highlighted in red. (B) Manually gated neutrophil subpopulations from Figure 3A were back gated to the viSNE map (right plot in each population). The automated clusters from Figure 1B are shown on the viSNE map (left plot in each population) to indicate the location of the corresponding automated gate. The percentage on the plots indicates how many cells of the manually gated neutrophil subpopulation fall into the automated gate. (C) The manual strategy to select blood neutrophils shown in supplemental Figure 4 were validated with flow cytometry.

**Supplemental Figure 6. Neutrophils display increased heterogeneity in melanoma-patient blood.** This supplemental figure is associated with Figure 3E. viSNE maps show FLOWSOM clusters from healthy donors’ blood neutrophils (right) compared to treatment-naïve melanoma patients (left). 2 healthy donors were analyzed.

**Supplemental Figure 7. The phagocytic and ROS-producing capacities of the neutrophil subpopulations compared to other leukocytes.** This supplemental figure is associated with Figure 4. (A) The percentage of the Zym+ cells in hNeP and N_1_ compared to that in Monocytes. Each dot represents result of one patient. (B) The geometric MFI of Zymosan in hNeP and N_1_ compared to that in monocytes. Error bars indicate mean with SD. (C) The frequency of Zym+ neutrophil subpopulations in total Zym+ neutrophils. (D) The frequency of Zym+ neutrophil subpopulations in total Zym+ leukocytes. (E) The percentage of the ROS+ cells in N6 compared that in monocytes and lymphocytes (T cells, B cells, and NK cells) from the + sti group. Each dot represents result of one patient. (F) Bar graphs show the efficacy of the ROSi in each gated neutrophil subpopulation. Error bars indicate mean with SD.

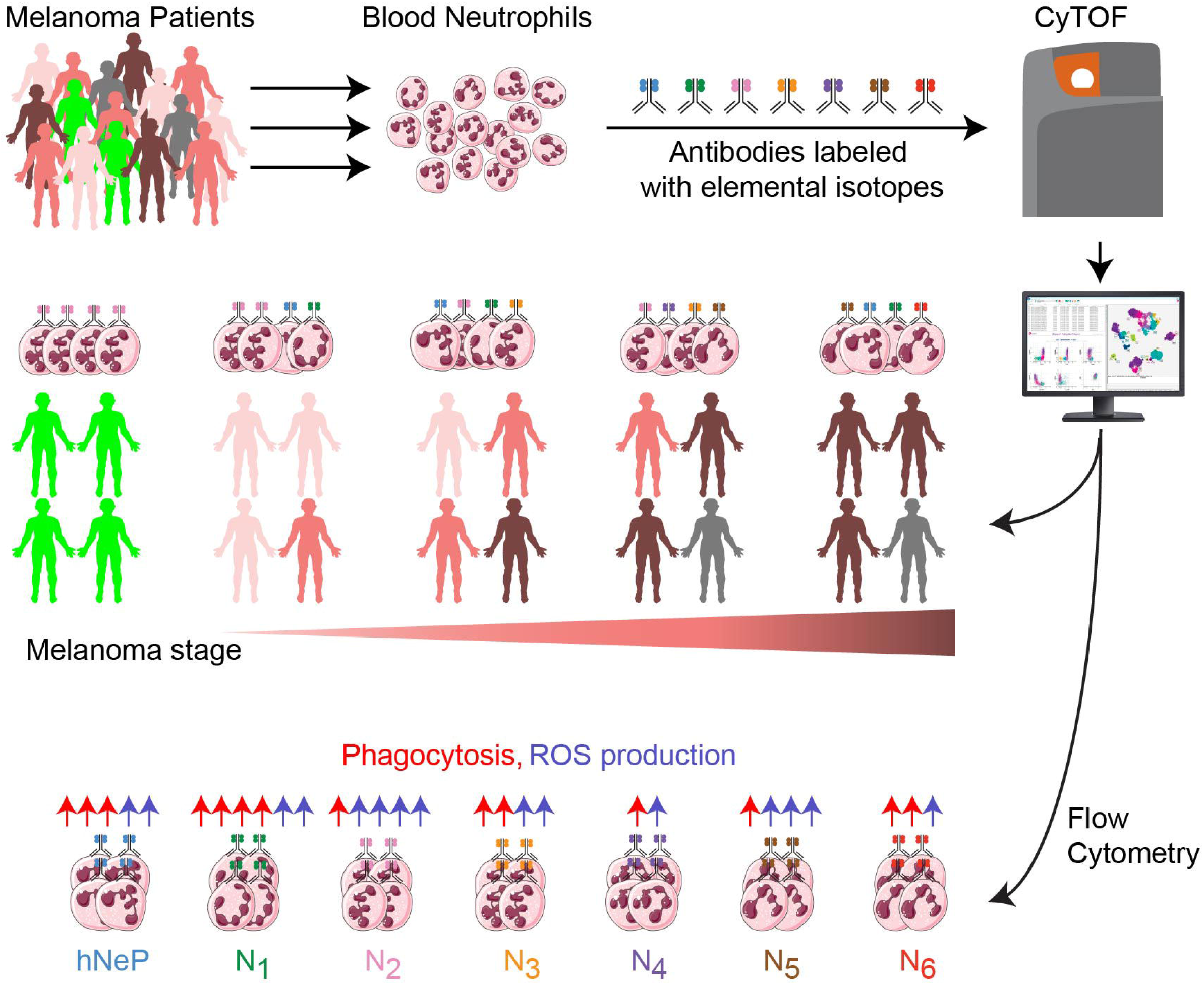

