## Supplemental Figures for "CyTOF reveals phenotypically-distinct human blood neutrophil populations differentially correlated with melanoma stage"

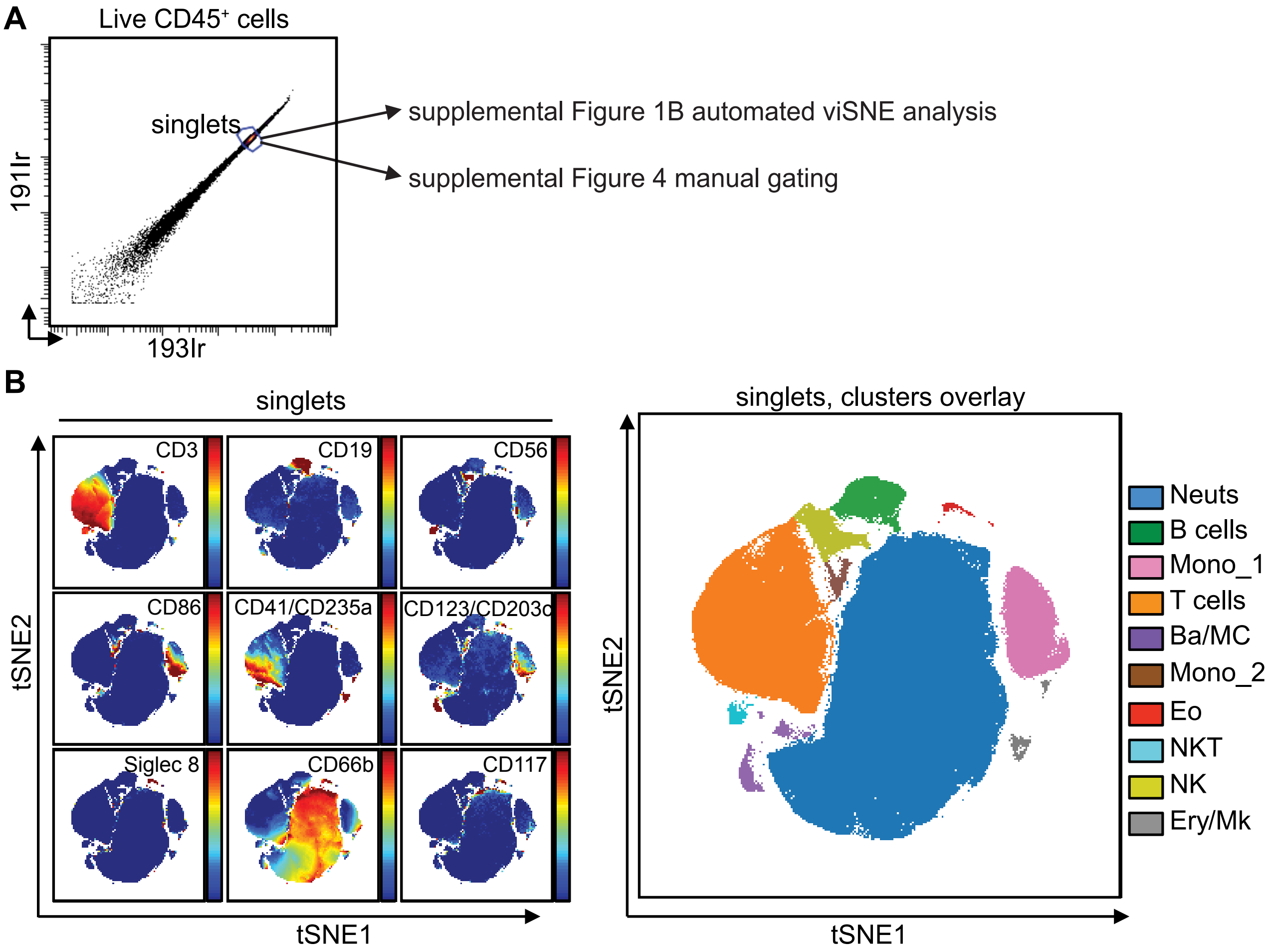


Supplemental Figure 1. Gating strategies to identify the CD66b+ blood neutrophil cluster in melanoma patients with CyTOF.

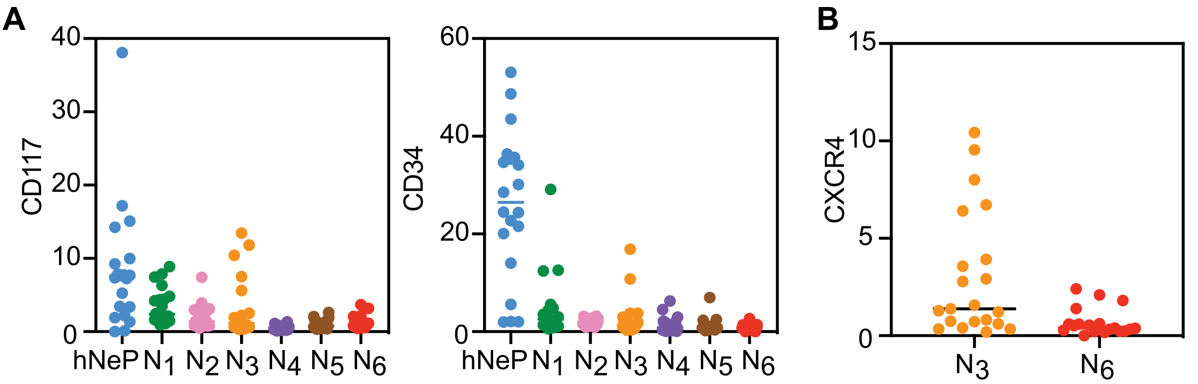


Supplemental Figure 2. Expression levels of markers in 7 automated blood neutrophil clusters.

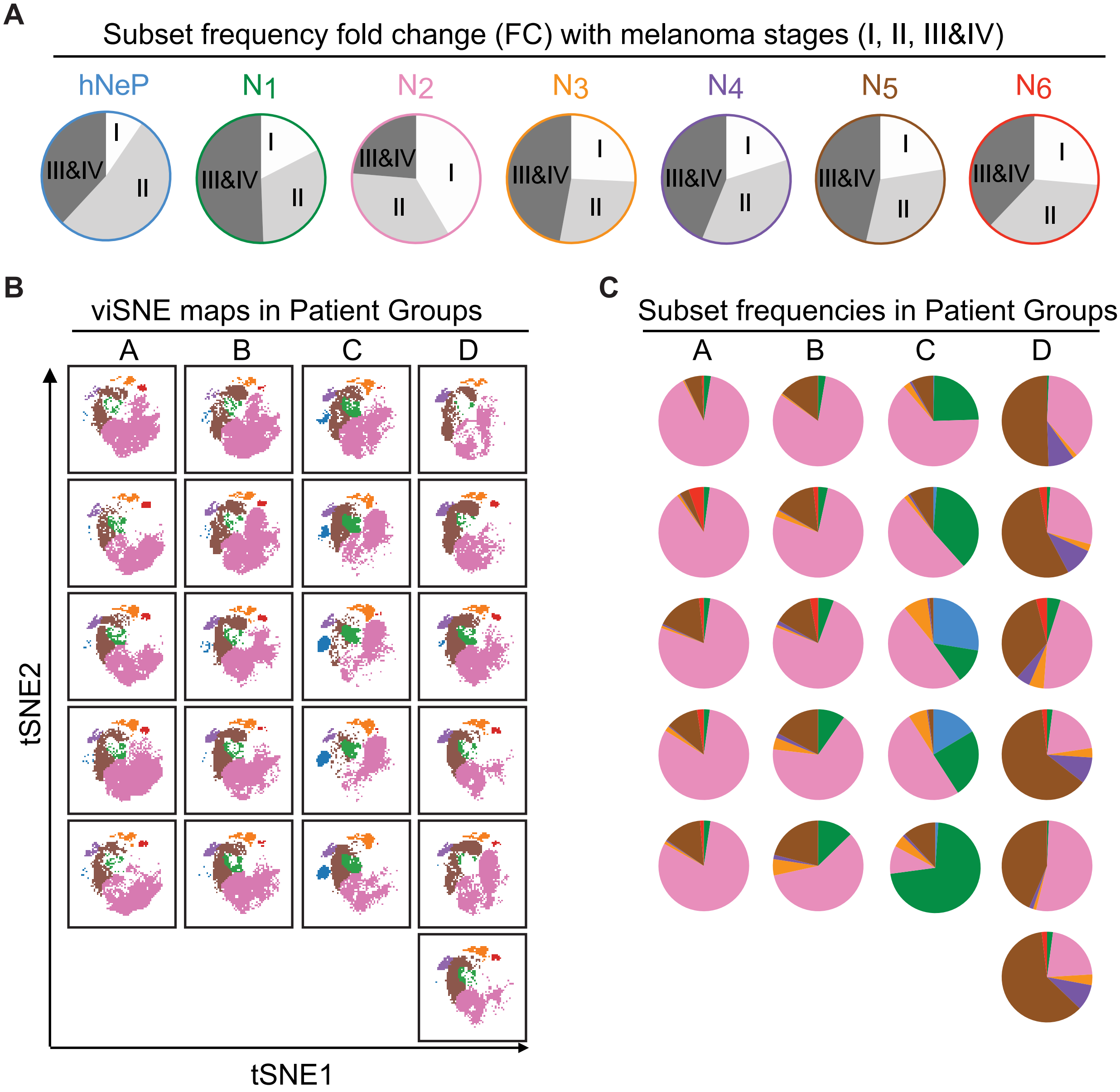


Supplemental Figure 3. FLOWSOM clusters (hNeP and $N_1$-$N_6$) shown on viSNE maps categorize melanoma patients into different groups.

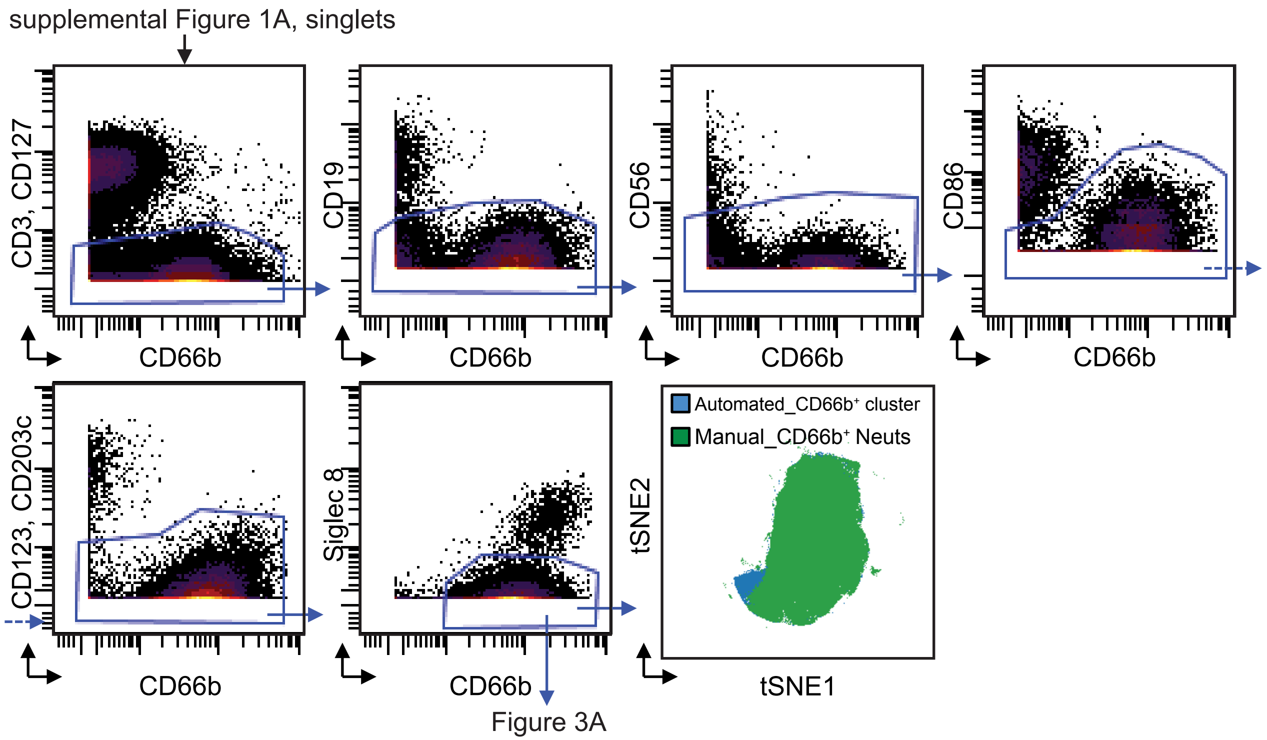


Supplemental Figure 4. Manual gating strategy to select CD66b⁺ blood neutrophils with CyTOF.

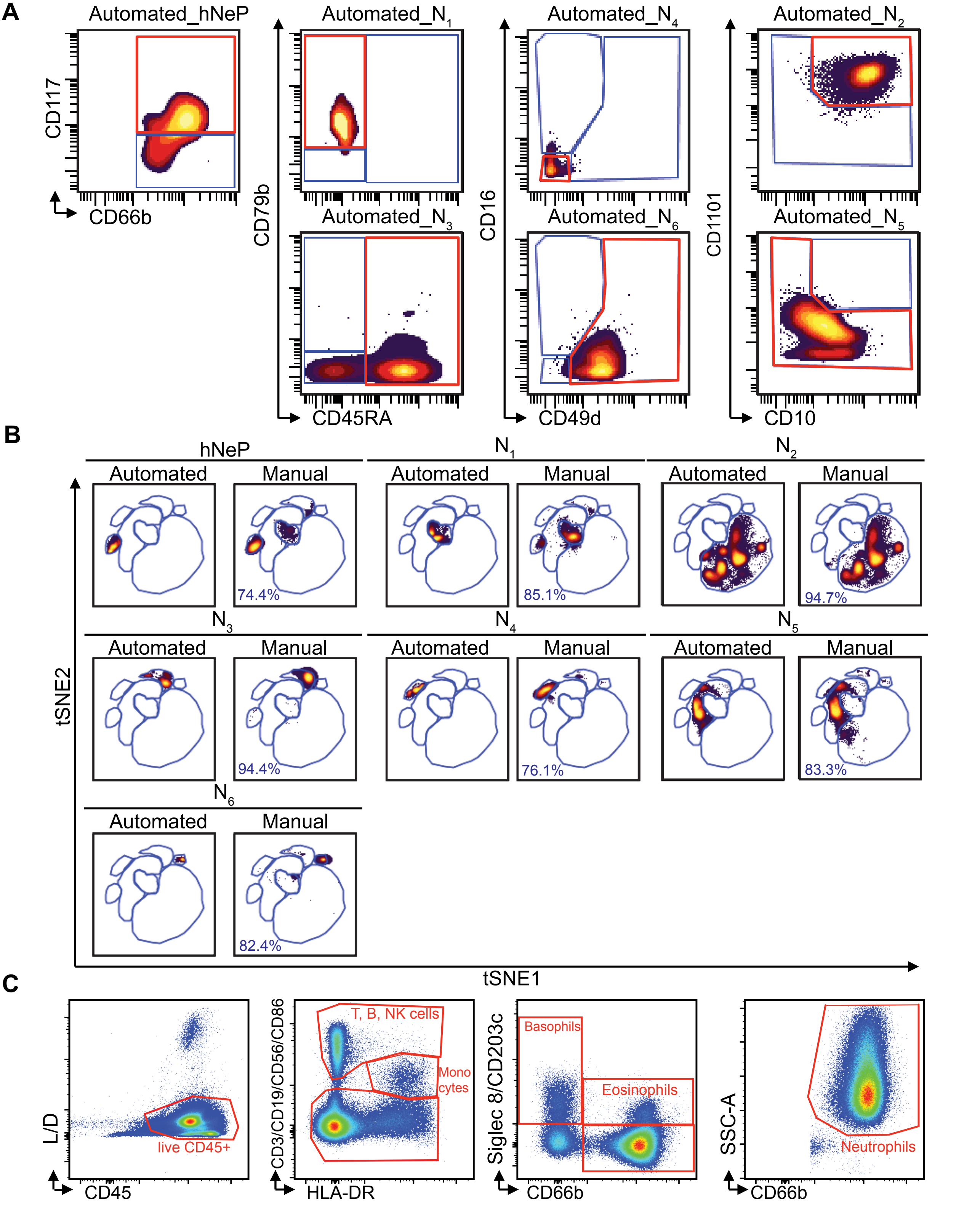


Supplemental Figure 5. Manual gating strategy recapitulates automated neutrophil clusters.

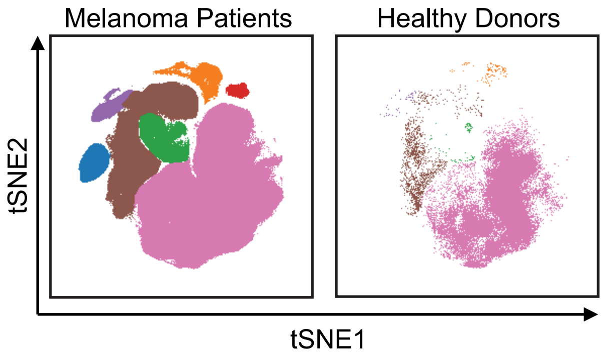


Supplemental Figure 6. Neutrophils display increased heterogeneity in melanoma-patient blood.

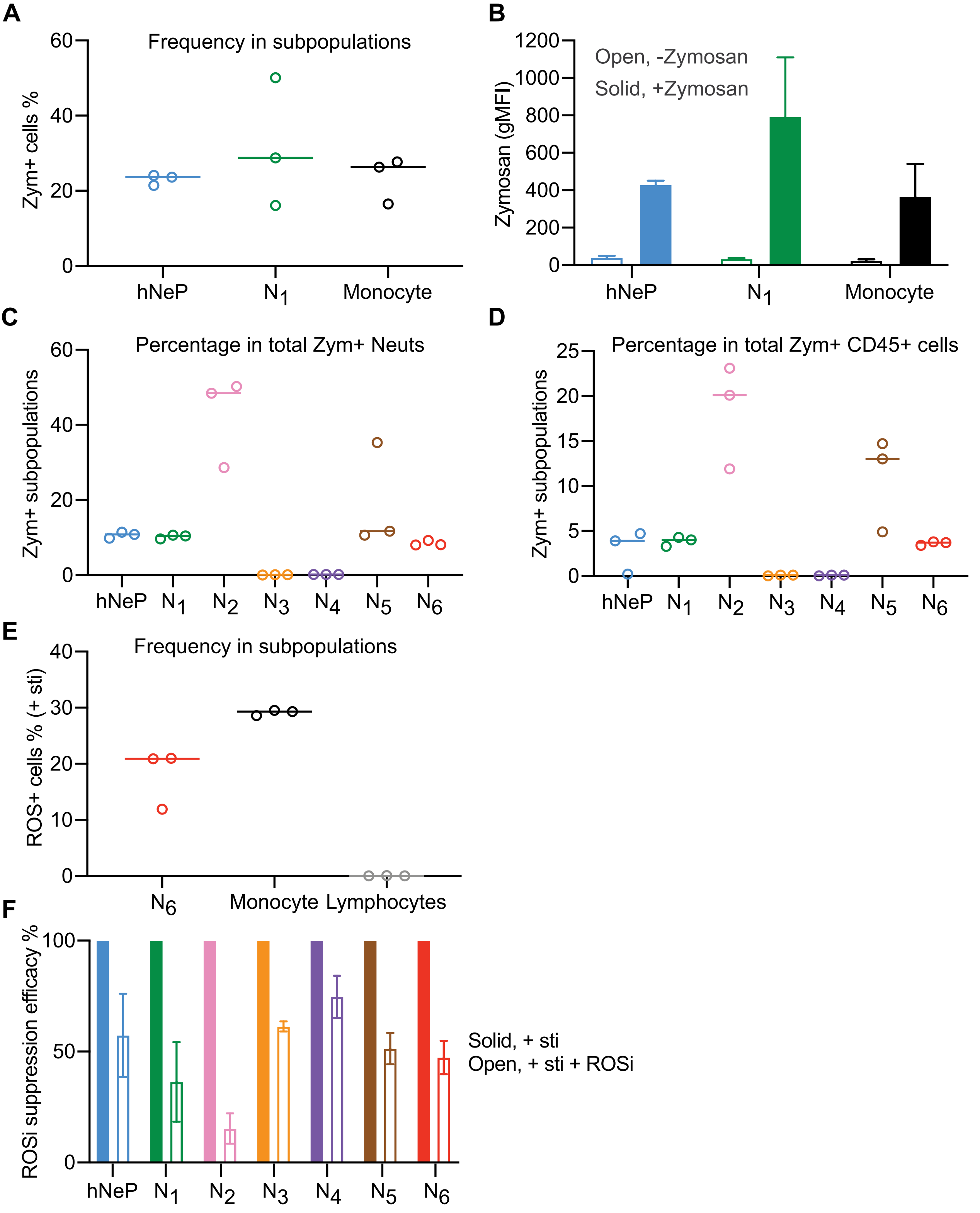


Supplemental Figure 7. The phagocytic and ROS-producing capacities of the neutrophil subpopulations compared to other leukocytes.
